# Trials and tribulations of Neotropical plant taxonomy: pace of tree species description

**DOI:** 10.1101/2023.09.05.556231

**Authors:** Manuel Luján, Rosalinda Medina Lemos, Eve Lucas, Fabián A. Michelangeli, Ghillean Prance, Terry D. Pennington, Jerzy Rzedowski, Daniel Santamaría-Aguilar, Ella Serpell, Cynthia Sothers, Alexandre R. Zuntini

**Affiliations:** Royal Botanic Gardens, Kew, Richmond, Surrey, UK; UNAM, Mexico; Institute of Systematic Botany, The New York Botanical Garden, Bronx, New York, USA; INECOL, Mexico; Louisiana State University, Baton Rouge, Louisiana, USA; Missouri Botanical Garden St. Louis, St. Louis, Missouri, USA

**Keywords:** fieldwork, herbarium, Latin America, Neotropics, new species, specimen collection, species description

## Abstract

One of the main aspects of taxonomic research is the description of new species. Identifying how to describe new species more efficiently is key to completing the inventory of the Neotropical flora in an era of massive biodiversity loss.
Here we calculate the interval between first specimen collection and new species publication for a group of 2126 Neotropical trees, and discuss the historical context surrounding specimen collection and new species publication events.
Our results reveal that on average, it takes almost 16 years from specimen collection to publication of a new Neotropical tree species, which is considerably shorter than previous estimates for other tropical groups. The central Andes is the region that had the longest average time lags, while the Choco had the shortest. Peru had the longest average time lags by country, while Haiti had the shortest. The average time lags increased until the early 1900s, when a decrease was observed, with the shortest lags between 1941 to 1960.
We found that the majority of the species described more rapidly are from plants collected and described by the same researcher. We demonstrate how political instability and conflict can delay or impede the completion of research initiatives in the region. We argue that enhancing international collaboration and training opportunities in Latin American countries, as well as ensuring safe plant collection campaigns, are critical to complete the inventory of the Neotropical flora.

**Societal Impact Statement:** This study aims to answer the question of how long it takes to describe a new species of tree in the Neotropics by calculating the time elapsed from collection of the first specimen to the publication of a new species. Given the current unprecedented concerns for global biodiversity loss, it is critical to identify best practices for describing new species more efficiently so we can complete the inventory of the Neotropical flora and implement appropriate conservation strategies in a timely manner.

## Introduction

Naming living organisms is a fundamental step in the process of inventorying, using, and conserving biological diversity. Throughout the history of biology, profound conceptual and technical changes have transformed the practice of identifying and naming species (Mishler, 2021 and references therein). Early species concepts used typological approaches that relied on key characters to identify species. Subsequent natural systems statistically analysed characters to group organisms based on numerical similarity (e. g. phenetic species concept, Sokal and Crovello, 1970), while others proposed using monophyly as main criterion to define species as clades (e. g. phylogenetic species concepts, Mishler and Donoghue, 1982; Mishler and Brandon, 1987). Modern approximations consider multiple properties such as diagnosability, monophyly and distribution, as lines of evidence that identify species as distinct lineages (unified species concept, de Queiroz, 2007). More recently, it has been suggested that species could be defined using heuristic (Wells *et al*., 2022) or pluralistic (Mishler, 2021) approaches, in which patterns of genetic, morphological, and ecological variation intrinsic to each group of organisms are assessed in a case by case scenario, without attempting to apply a universal set of criteria to define all species. Using multiple lines of evidence has contributed to consider taxonomy not only as the practice of naming and classifying organisms, but also as a bona fide scientific discipline in which species are testable and falsifiable hypotheses. Regardless of the prevailing conceptual framework, describing and delimiting species has been, and continues to be, a scientific process that requires an appropriate level of taxonomic capacity, and more relevant for the present work, time.

The average time from the first collection of a specimen to its formal description and naming as a new species has been estimated to be 21 years (Fontaine *et al*., 2012). In plants, this rate has been estimated to be almost 39 years, although for tropical groups the process may take slightly longer (e. g. 41 years in the genus *Aframomum* [Goodwin *et al*., 2020]). In addition, the interval between specimen collection and description of a new species of flowering plant seems to have stagnated since the 1970’s (Bebber *et al*., 2014). The evident lag between specimen collection and species description is part of the taxonomic impediment, a concept coined by Taylor (1983) to explain the disparity between our limited taxonomic capacity and the full magnitude of biodiversity yet to be described.

Biodiversity is unevenly distributed across the world, with the Neotropical region having the highest diversity of plant and animal species (Raven *et al*., 2020). Out of the 383,671 accepted species of vascular plants (Lughadha *et al*., 2016), the Neotropics houses the highest levels of species richness comprising around 90,000–110,000 plant species (Antonelli and Sanmartín, 2011; Govaerts, 2001), of which 23,631 species can be classified as trees. An estimated 30% of all tree species are at risk of extinction, and the Neotropics are one of the regions where more trees are threatened with over 8,000 species at risk (Botanic Gardens Conservation International, 2021).

Given unprecedented concern for species extinction and conservation, here we aim to better understand a key aspect of the taxonomic impediment in plants. In this study we revisit the question of how long it takes to describe plant species, specifically focusing on Neotropical trees. We aim to estimate the time elapsed between specimen collection and new species description in key groups of ecologically important tree lineages occupying a wide diversity of tropical habitats. We discuss our results in the context of the historical and current conditions influencing taxonomic capacity in the Neotropical region, thus identifying opportunities to shorten the period between collection and publication and accelerating output of new biodiversity data.

## Materials and Methods

We selected groups of trees that are ecologically important across the Neotropical region in terms of species richness or relatively high local abundance, and for which sufficient taxonomic expertise allows efficient verification of type specimen information. We used the standard concept for a tree defined as a perennial woody plant with secondary thickening and a clear main trunk (Beentje, 2020). The groups selected include species in *Bursera* (Burseraceae), *Clusia* (Clusiaceae), *Myrcia* sect. *Calyptranthes* (Myrtaceae), *Inga* (Fabaceae), *Virola* (Myristicaceae), as well as tree species in Bignoniaceae, Chrysobalanaceae, Humiriaceae, Lecythidaceae, Melastomataceae, Meliaceae and Sapotaceae (for the complete list of species see Appendix 1). We included species that commonly grow as trees as well as some species that grow as woody hemiepiphytic plants that can also develop into trees. We used the World Checklist of Vascular Plants (Govaerts *et al*., 2021) to record the date of publication for each accepted species. Then, we recorded the collection date of the earliest plant material cited in each species’ protologue, including (but not restricted to) holotypes, syntypes, and paratypes. When collection dates were recorded as a range, we chose a conservative approach and selected the most recent year as collection dates. In cases where collections dates were not recorded, these were estimated based on the collector’s biography, expedition’s account, or other historical source of information (e. g. Prance, 1972 a). For species originally described under different names, we used the information of the earliest basionym. Species were excluded from the analysis if accurate dates were not available for the plant material used in the original description. For each species included in the analysis we calculated the interval between the date of the earliest collection and the date of effective publication. We classified type locality of each species by country and by region according to the Terrestrial Ecoregions of the World from the World Wildlife Foundation (WWF; Olson *et al*., 2001), with modifications for the Neotropics by Freitas *et al* (2019). We calculated average lags from first specimen collection to new species publication by country, ecoregion, and within 20-years intervals. We compared average distributions using Kruskal-Wallis H tests and post-hoc Dunn’s test using the Bonferroni correction. We assembled timelines of specimen collection and species publications by country and by region where type material was collected, and identified publications where highest numbers of tree species have been described.

Finally, we described some of the social and political events that may have influenced the conditions for doing field expeditions to collect plant material in the Neotropical region as well as the publication of new species.

## Results

### Time lags between specimen collection and new species description

From a total of 2126 Neotropical tree species studied, the average time lag between first specimen collection and publication of a new species was 15.7 years (± 20.4 SD). Half of the species evaluated in this study were described 4 – 20 years after the first specimen collection. One percent of the species (21 spp) have time lags longer than 100 years, and 1.5% (30 spp) have time lags shorter than one year. The longest time lags are recorded in two species of Melastomataceae, *Axinaea confusa* E. Cotton from Peru (229 years) and *Meriania aguaditensis* H. Mend. & Fern. Alonso from Colombia (204 years).

The longest average time lags by ecoregion were recorded for species described from the Caribbean dry forest (26.5 years ± 33.3, n = 4) and the central Andes (23.4 years ± 30.1, n = 102), while the shortest average time lags were recorded from the Choco (9.5 years ± 11.5, n = 113), and xeric Mesoamerica (11.2 years ± 12, n = 17 [Fig. 1]). The Kruskal-Wallis H test indicates the differences between time lags across ecoregions are statistically significant (H[14] = 76.5, *p* < 0.001).

**Figure 1.**
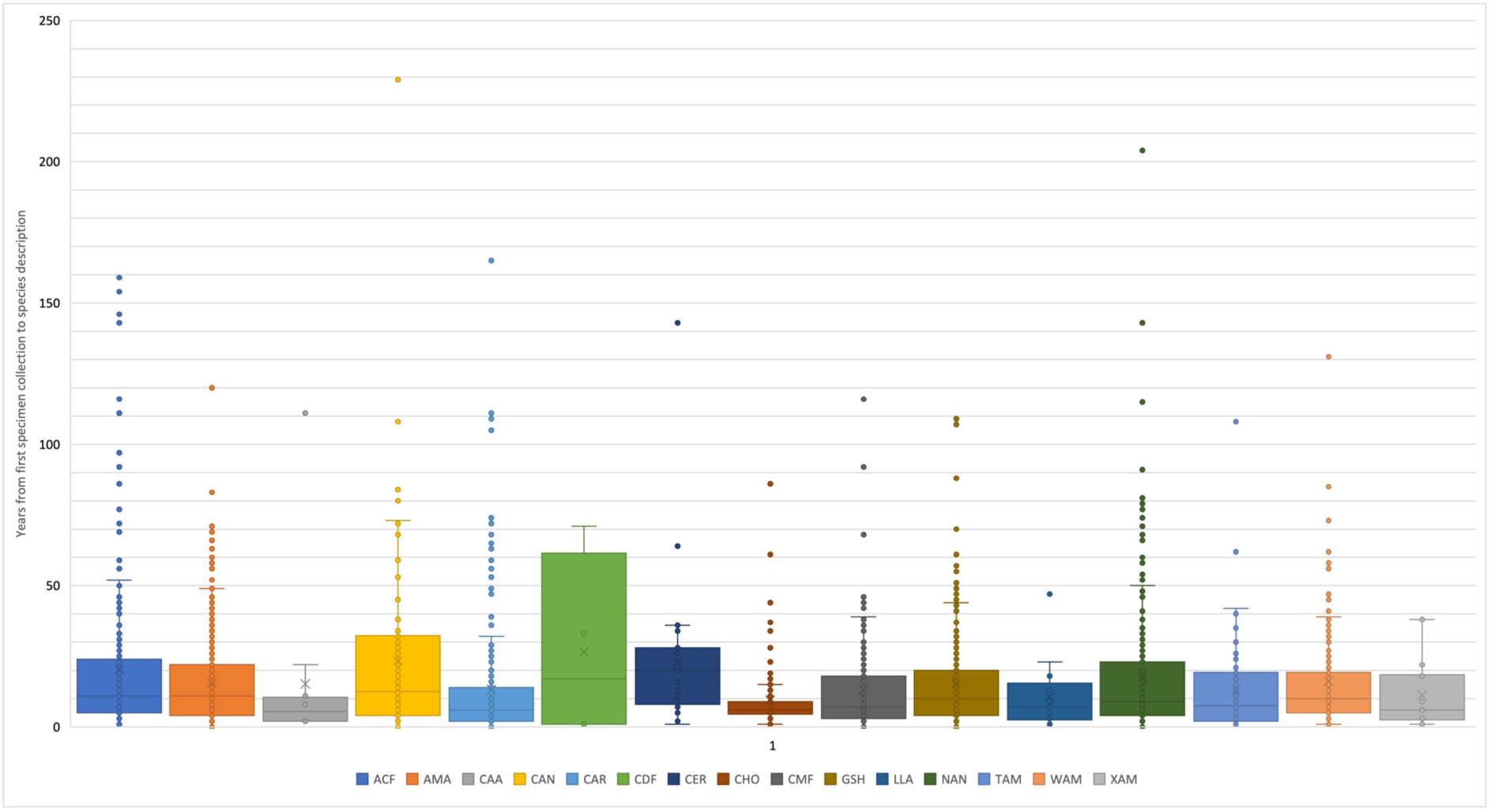
Time lag between oldest collection and new species description, summarised by Biome. Biomes code follow Freitas *et al* (2019): ACF: Atlantic Coastal Forest; AMA: Amazon; CMF: Central American Moist Forest; CAA: Caatinga; CAR: Caribbean; CDF: Caribbean Dry Forest; CAN: Central Andes; CER: Cerrado; CHE: Chaco and 750 Espinal; CHO: Choco; GRP: Grassland and Pampa; GSH: Guiana Shield; IAF: Inter751 Andean Dry Forest; LLA: Llanos; NAN: Northern Andes; PAN: Pantanal; SEU: Southeastern United States; TAM: Tropical Central American Dry Forest; WAM: Western Amazon; XMA: Xeric Mesoamerica.

The longest average time lags by country were recorded from Peru (20.6 years ± 25.2, n = 182) and Jamaica (19.1 years ± 29.9, n = 29), while the shortest lags were observed in Haiti (4.3 years ± 6.1, n = 23) and Belize (5.1 years ± 8, n = 12). Of the countries with more than 100 species included in this study (Brazil [579], Colombia [265], Cuba [112], Ecuador [127], Mexico [129], Peru [182] and Venezuela [175]), the longest time lags were observed in Peru and Brazil (17.2 years ± 21.3), while the shortest time lags were recorded in Cuba (12.8 years ± 17) and Mexico (13.6 years ± 14.7 [Fig. 2]). Differences between time lags across countries are statistically significant (H[18] = 81.8, *p* < 0.001). At a regional scale, the longest collection to description lags were recorded in the Caribbean dry forest (CDF) and Central Andes (CAN).

**Figure 2.**
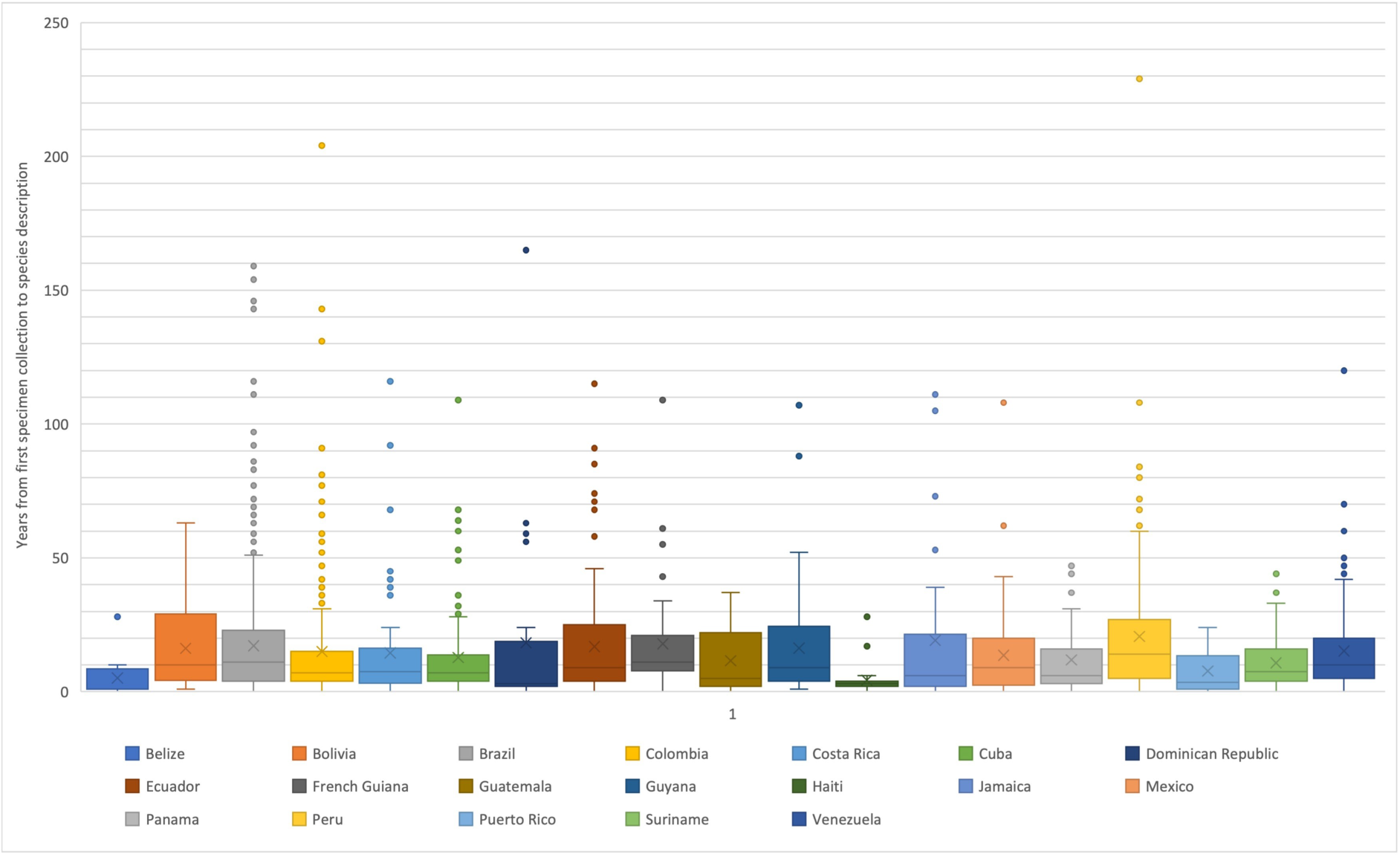
Time lag between oldest collection and new species description of Neotropical trees in our sample, summarised by country.

Average time lags increased from the beginning of records until the early 1900s, when a sudden decrease can be observed (Fig. 3). The shortest time lags are registered between 1900 and 1960, particularly between 1941 to 1960, when species were described relatively shortly after specimens were collected (8.7 years lag on average). After that, the average time lag increased significantly, the longest observed being since the year 2000 until the present (26.4 years on average for species described in 2001 – 2020 [Fig. 3]). Differences between time lags across the 20-year periods are statistically significant (H[12] = 250.13, *p* < 0.001).

**Figure 3.**
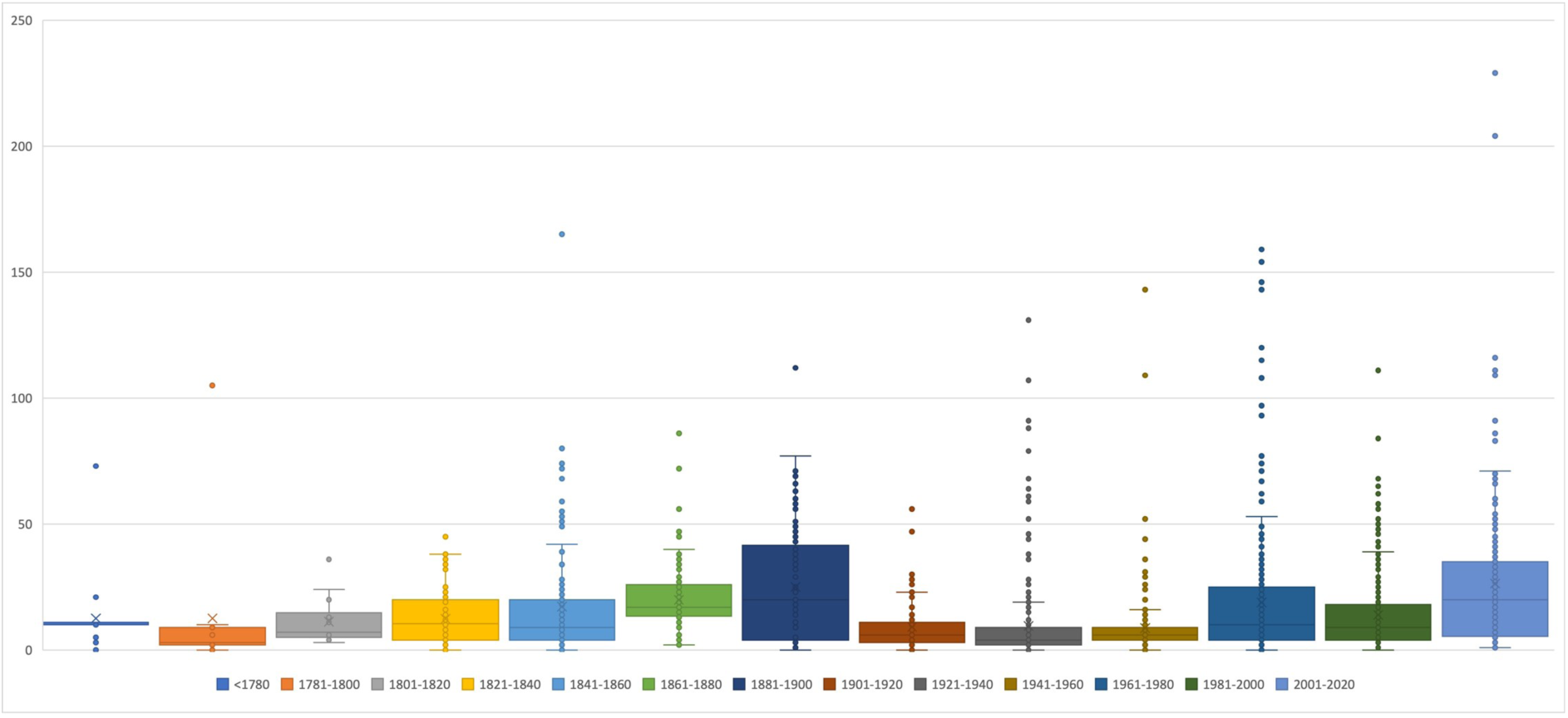
Time lag between oldest collection and species description of Neotropical trees in our sample, distributed in 20-years intervals.

A timeline of the total number of new species descriptions shows that from 1800 to 1860 there was a stable increase in the number of new species described per period, followed by a slight decrease until the early 1900’s (Fig. 4). From 1920 onwards there was a steep increase in the number of new species described per period that reached its peak between 1960 and 1980, when 340 species of Neotropical trees from our sample were described. The number of new species described per period shows a significant decline from the beginning of the year 2000 until present (Fig. 4).

**Figure 4.**
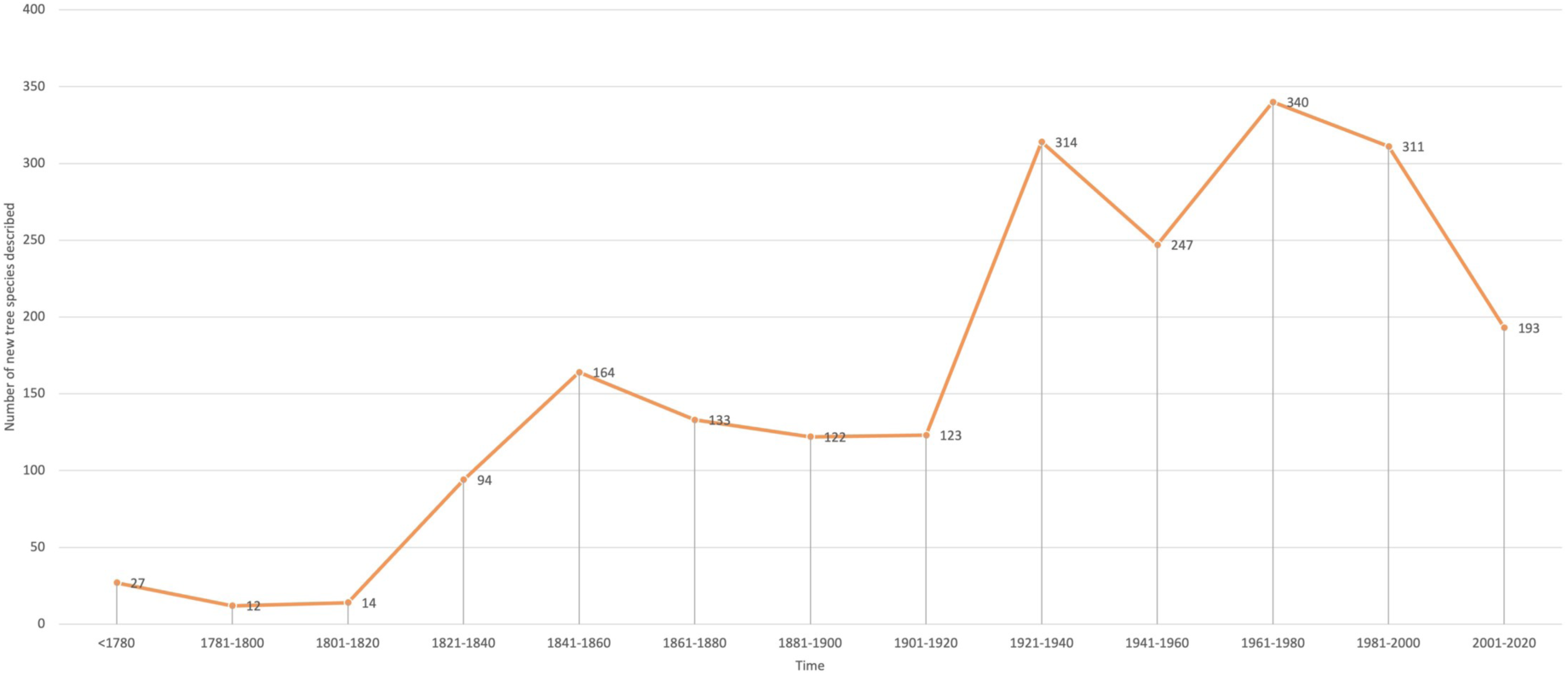
Number of species described through time. Time is divided in 20-years periods. Number of new tree species described since 2020 in our sample is 108, although this period is not depicted.

The period between 1900 and 1960 shows the shortest time lag from collection to description of new tree species in our data for the region. After 1960, the time lag from collection to description of new tree species increased significantly. Notably, the period from 1960 to 1981 shows an average of 18.9 years time lag, more than double than the average of the three precedent periods (9 years). Similarly, the period from 2000 to 2021 shows an average 26.4 years time lag for a new species to be described, the longest recorded for any period in our records.

### On the origin of specimens and publication of new species

The earliest collections of plant material used for describing new species of Neotropical trees included in our study date from Jamaica in the late 1600’s, the oldest collections of type material are of *Clusia flava* Jacq. collected in 1687 and described in 1760; and *Bursera simaruba* (L.) Sarg. collected in 1688 and described in 1753. A significant number of the earliest plant collections used to describe new species of Neotropical trees included in our study are from territories currently known as French Guiana, Guyana and Suriname [Figs. 5A and 5E]). Over two thirds of the tree species described before 1800 were described by Jean Baptiste Aublet, who collected in French Guiana from 1762 to 1764, and described over 400 new plant species (Howard, 1983). On average, Aublet species were described 11 years after collection. Since the late 1700s until now, most type collections originate from Brazil, with an important number of collections from Colombia, Ecuador, Mexico, Peru, and Venezuela (Fig. 5B-D). Notably, during the period of 1941–1960, the largest number of newly described species were from Colombia (Fig. 5F-H). Timelines of the number of new species described per year for countries with over 100 species included in this study allow us to identify the publications in which most Neotropical tree species included in our sample have been described (Figs. 6A-G).

**Figure 5.**
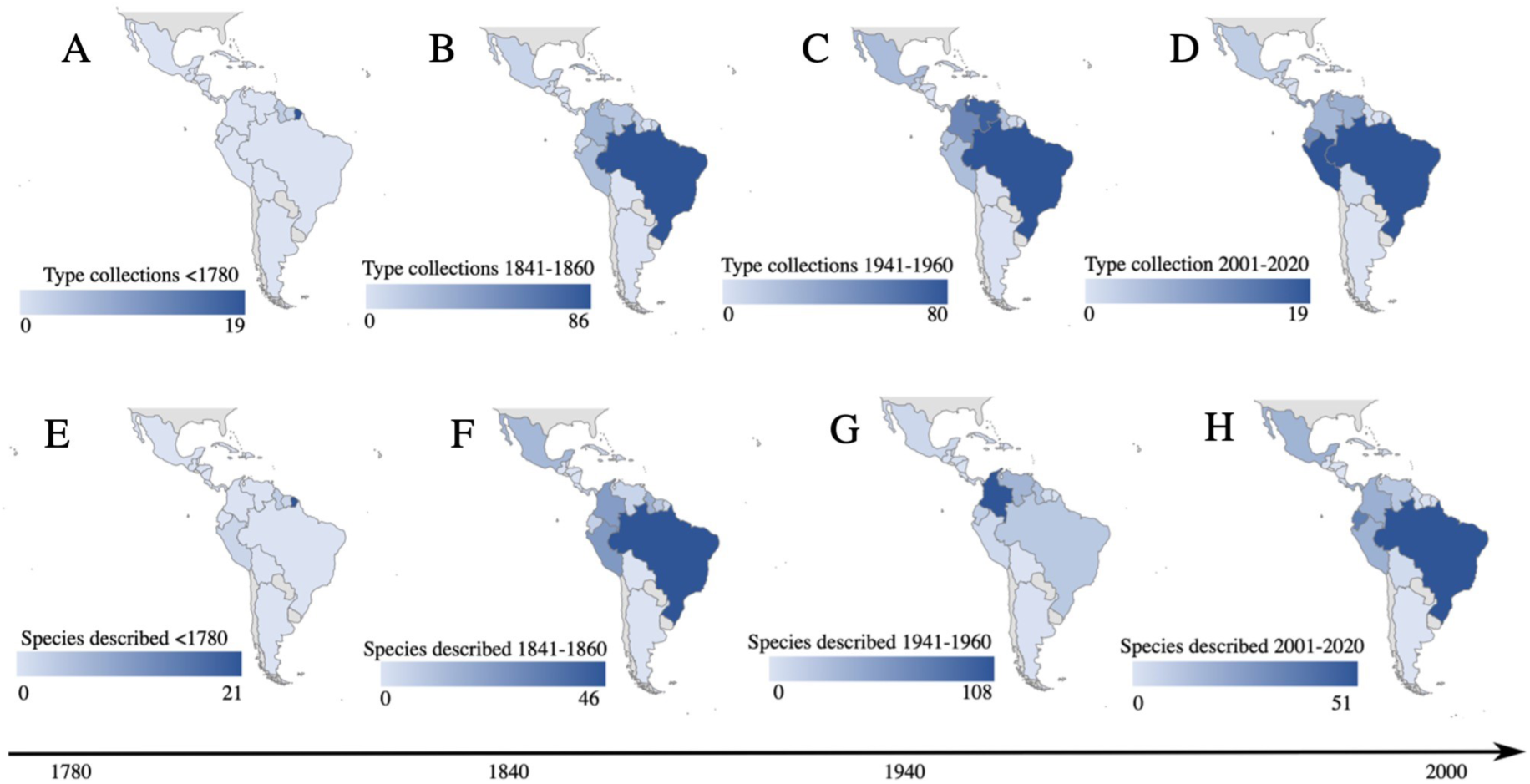
Number of collections of type specimens (A-D) and new tree species described (EH) by country and time periods.

**Figure 6.**
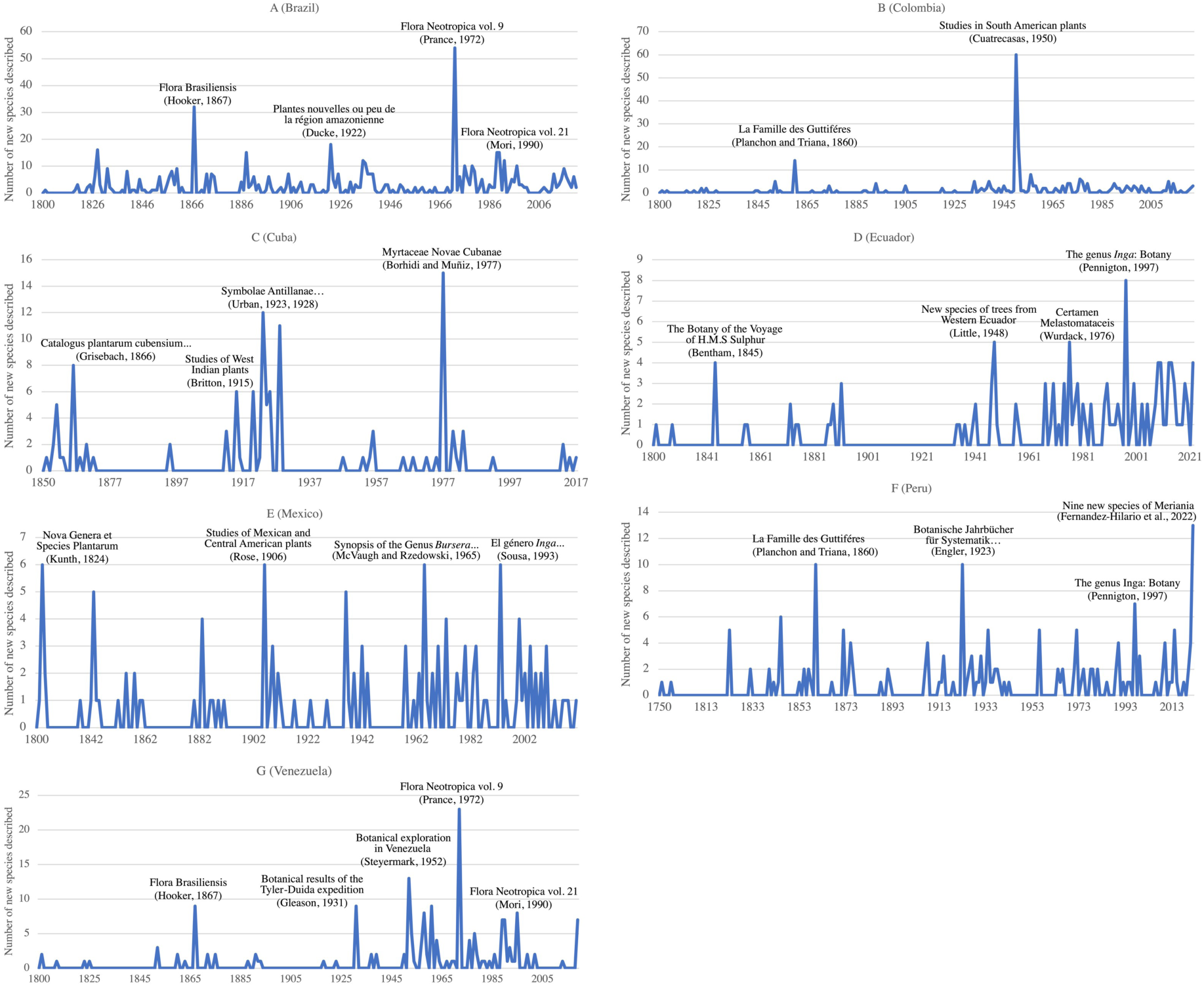
Number of new species described per year for the six countries with >100 species of trees in our data. Key publications related to description peaks are highlighted.

## Discussion

We analysed the pace of Neotropical tree species description considering the time, place and historical context that may have influenced plant collection and new species publication events. The average time (almost 16 years) between specimen collection and publication of our sample of Neotropical trees is considerably shorter than estimates in other tropical groups such as the understory genus *Aframomum* (41 years [Goodwin *et al*., 2020]). It stands to reason that more interest has been given to documenting, describing, and managing species of trees because they are prominent components of terrestrial ecosystems that are sources of several natural products such as timber, food, resins, etc., as well as provide critical ecosystem services (Boyd *et al*., 2013). Future studies might test whether the collection to description lag varies according to plant habit, as plants with life forms that make them relatively more difficult to collect, identify and preserve, may have longer species collection to description lags (e. g. aquatic plants [Bedoya and Olmstead, 2022]).

### Time lags by region and country

The extended time lag observed in the Caribbean dry forest (26.5 years average) is based on a relatively small group of samples (n = 4) that is strongly biased by the long time lag (71 years) of *Handroanthus billbergii* (Bureau & K.Schum.) S. O. Grose. The average of the other 3 species recorded from CDF is 11.6 years, below the global average of all species in this study. The Central Andes is the region with over 100 species sampled in this study that shows the longest collection to description time lag. Similarly, the longest time lag by country is recorded for Peru, a country that occupies most of the Central Andes’s region. Botanical exploration in the Central Andes, particularly in Peru, started ca. 1750 with de Joseph de Jussieu who collected plants as part of a scientific expedition originally sent by the French Academy of Sciences, although little is documented from his field campaigns. During the late 1700s, Hipólito Ruiz and José Antonio Pavón collected extensively in the Central Andes region as part of the Botanical Expedition to the Viceroyalty of Peru commissioned by King Charles III of Spain, however much of the plant materials collected during these early expeditions were lost in fires and shipwrecks, and it was not until 1798 when some new species were finally published by these authors. Due to high publication costs, and instability caused by the Napoleonic wars, Ruiz and Pavón were unable to publish most of the findings of their botanical explorations in the Central Andes (Weberbauer, 1911). It was in 1860 when many Ruiz and Pavón plant materials and collection data were finally used to describe and publish new species, particularly by Jules E. Planchon and José J. Triana. Observing the top ten plant species with the longest time lags from Peru, six of them are based on Ruiz and Pavón Andean specimens that were described by Planchon and Triana, over 72 years after collection. This clearly illustrates how the completion of scientific endeavours can be delayed due to the difficult conditions under which specimens are often collected in nature, as well as by political instability and conflict. Also, how more than one generation of researchers may be involved in seeing material from collection to description. In contrast, the shortest time lag recorded by region, from the Choco (Fig. 1), is explicable due to the high numbers of species collected and described by the same researcher. For example, Alwyn H. Gentry collected and described several new species of Bignoniaceae from the Choco region himself (e. g. Gentry [1973]). Likewise, Jose Cuatrecasas collected extensively in the Choco and described several new species (e. g. Cuatrecasas [1950]). In fact, most of the species with the shortest time lag from the Choco are from Cuatrecasas, who on average described new Chocoan tree species in less than six years after specimen collection. Cuatrecasas’ prolific scientific production led him to not only describe new species fast, but also in large numbers. Colombia was the country with the highest number of tree species described during 1941–1960 mainly due to Cuatrecasas’ publications (Fig. 5G). Most of Cuatrecasas’ species of trees from the Choco are from plants he collected from 1942 to 1947 while he was based in Cali as the director of the the department’s official botanical commission. In 1947 he took a position as Curator of Botany at the Field Museum of Natural History in Chicago, from where he described several new tree species based on his collections from the Choco (Huber, 1996). After this highly productive period in terms of plant collections and publication of taxonomic novelties, botanical field research likely declined in some regions in Colombia during the early 1960s when two main guerrillas emerged, the ELN (Spanish acronym), active in the valley of the Magdalena River and FARC (Spanish acronym), present in various regions (LeGrand, 2003). During early stages of the conflict, areas controlled by the guerrillas were bombed by the military, probably deterring scientists from undertaking fieldwork in some areas. Our data show that in 1964 and 1965, when the FARC were formed, no type material used to describe new tree species was collected in Colombia, compared to neighbouring Ecuador and Venezuela where plant collections that rendered type specimens continued to be made (5 and 2 type collections during the same period respectively). Since the guerrillas started, scientific exploration in some regions of Colombia has been challenging due to security concerns, although a peace deal between the government and FARC signed in 2016 led to a revamp in field research (Wade, 2018; Irwin, 2023), and several expeditions have been undertaken (e. g. Lucas *et al*., 2023). More plants are being collected from poorly documented areas, and it is expected that new Colombian plant species will be described in the forthcoming years.

The shortest collection to description lag for Haitian species is likely due to the collections of Erik L. Ekman, published by Ignatz Urban as part of a prolific collaboration to document a significant part of the Antillean flora. Ekman made extensive collections in Haiti from 1924– 1928 supported by Urban, who received Ekman’s collections in Berlin and published extensively (Standley, 1931). The average time lag for Ekman-Urban tree species in Haiti is 4.2 years. Evidently, this was an efficient model for describing new plant species, allowing a field botanist to collect year-round in an area, and send materials to a herbarium botanist with readily access to a large plant collection (Berlin herbarium before WWII [Hiepko, 1987]) as well as the resources to publish their results effectively. Admittedly, this approach could be interpreted as an unethical practice where scientists from affluent countries, active in less developed ones, source research materials without necessarily involving local partners in the project design and publication of results (Baber, 2016; Li, 2021). Since 1993, The Convention on Biological Diversity (CBD, 1992), and its supplementary agreement the Nagoya Protocol (NP, 2011) were signed to articulate international efforts to ensure fair and equitable sharing of benefits arising from the utilisation of biodiversity and its genetic resources. United Nations’ member states committed to these multilateral agreements developed national strategies to regulate access to genetic resources and traditional knowledge including prior informed consent of the party providing resources, sharing research results and promoting technical and scientific cooperation. Most international scientific botanical institutions currently comply with CBD and NP obligations, ensuring botanical research is now carried out in a more fair and equitable way.

In Belize and Puerto Rico, most tree species included in this study follow a similar pattern as in the Choco, where specimen collection and species description were carried out by the same person. In Belize new tree species were described by Paul C. Standley on average one year after collection, and in Puerto Rico, new tree species were described by Henri Alain approximately six years after collection (see Appendix 1).

Within the most rapidly described species from Cuba, are those published by August Grisebach (1866) based on materials collected by Charles Wright, who botanically explored the island between 1856 and 1867 (Stearn, 1965), as well as species described by Nathaniel L. Britton and Elizabeth G. Britton, who published several new species from Cuba and other Caribbean islands based on specimens collected by them and by collaborators. Botanical exploration in Cuba, particularly by American institutions, diminished when the Cuban revolution started (i. e. no type collections for new tree species were made between 1957 and 1959, when the revolution took power), and ceased when the economic embargo was imposed by the USA in 1961 (i. e. no type collections for new tree species were made between 1961 and 1966). The embargo effectively blocked any American research funding from reaching Cuban institutions and restrained scientific collaboration between the two countries (Ronda-Pupo, 2021). Most of the new species of trees described from Cuba during the revolution were made by botanists associated to European institutions who explored the island during the mid 1900s, in particular Attila Borhidi (Hungary) and Johannes Bisse (Germany), who on average described new Cuban tree species 14.2 and 10.4 years after collection respectively. A window of opportunity for scientific collaboration between American and Cuban botanic institutions opened in 2015 when diplomatic ties between the USA and Cuba were re-established, thereby promoting the expansion of scientific collaboration. As an example, The New York Botanical Garden and Cuba’s National Botanical Garden launched new collaborative conservation research projects in 2015. However, US policies toward Cuba were revised in 2017, reversing some of the efforts made towards normalising diplomatic relationships. This illustrates how frequent policy shifts make research planning uncertain and may deter international scientific collaboration even in less restrictive times (Ronda-Pupo, 2021).

In Mexico, tree species described within one year or less after first specimens’ collection come from Cyrus L. Lundell and Faustino Miranda, who independently undertook botanical expeditions in Southern Mexico during the late 1930s, and rapidly published their own observations, including several new species descriptions (Rzedowski, 1967). According to our data, the period from 1940 to 2000 was the most prolific for botanical exploration in Mexico as type material for 63 new tree species were collected during that interval, the highest number of newly described tree species per period in our records. During these decades, several herbaria and research institutes were established, and a great number of Mexican botanists undertook extensive vegetation surveys across the country (Rzedowski, 1981). Beginning in the 2000s, particularly between 2006 to 2008, the Mexican government launched the “war on drugs’’, a national offensive against drug trafficking in which the military became involved in an effort to decimate drug cartels. Violence during this period was exacerbated when gun battles between security forces and cartel gunmen became increasingly common and deadly across the country (Grillo, 2013). This situation may have affected field scientific expeditions in Mexico since 2000. For instance, most species analysed from Mexico in our data are trees in the genus *Bursera*, collected and described by Jerzy Rzedowski between the 1960s and 1980s mainly from dry and seasonally dry forests. Areas where Rzedowski and collaborators first made their collections are now unsafe for other researchers to revisit due to the active presence of drug cartels, impeding thorough botanical documentation and delaying potential new species descriptions. A number of undescribed species of *Bursera* have not been published yet because plant materials needed for completing morphological descriptions such as mature leaves, female and male flowers, are lacking, and access to field sites is restricted. The number of type materials of new species of trees collected since 2000 in Mexico is four, one of the lowest recorded in our data. As a comparison, 21 new tree species have been collected in Brazil, 11 in Colombia, 36 in Peru since 2000. After decades of active botanical exploration in Mexico, it is possible that there may not be a great number of new tree species left to be described. However, we expect that safer access to field sites, particularly in poorly documented areas, will allow collecting plant materials that may render taxonomic novelties, not only in Mexico but in the entire region, as the Neotropics are predicted to contain over 30% of the world’s undescribed species of flowering plants (Joppa *et al*., 2011).

Regarding the high numbers of type collections and new species published from Brazil since 1800, this trend was particularly pronounced between 1961-1980, when 91 type collections were made and 96 new species described, the highest numbers recorded in Brazil and the entire region. It is interesting to note that publication of a series on Neotropical flora began in 1967 in New York Botanical Garden (NYBG) and continued actively through this period producing monographs of several woody plant families (Cowan, 1967). According to our data, the highest number of plant collections from Brazil during that period come from John J. Wurdack, Scott A. Mori and Ghillean T. Prance, who were based at NYBG and developed an active program of field research in the Amazonian and Guyanese regions contributing important volumes pf publications. During the period from 1961 to 1980 new Brazilian species were described 13 years on average after collection.

### Time lags by period

From the early 1800s until 1900 the collection to description time lag for tree species increased steadily reaching its peak between 1880 to 1900. Among the species with longest collection to description time lag are several in the genera *Clusia* (Clusiaceae) and *Licania* (Chrysobalanaceae), published by Adolf Engler (1888) and Karl Fritsch (1889) respectively. Species collected and described during this relatively slow period for taxonomic novelties were almost exclusively from continental America, mainly from Brazil. New species described by Engler were published over 40 years after collection on average, and were based on collections made by Richard Spruce, who explored the Amazonian region from 1849 to 1864 (mostly in Brazil, Ecuador, and Peru) and collected a vast amount of plant materials that served to described several new species (Seaward, 2010). Similarly, new South American tree species described by Fritsch were based on materials collected by Robert H. Schomburgk who explored British Guiana between 1835 and 1839 (Schomburgk, 1931). The average collection to description time lag for Fritsch-Schomburgk new tree species is 43 years.

Most of the species published between 1900 and 1960 are from Brazil (90), 48 of them collected and described by Adolpho Ducke, who extensively explored the Northern Amazonian region (Archer, 1962) and published new species of trees on average 6.6 years after collection. A significant number of species described during the first half of the 20th century, when new tree species were described relatively fast, come from Cuba (41 spp) mainly published by Urban, and from Peru (34 spp) mostly published by James F. Macbride who collected in Peru between 1922 and 1923, and published several new tree species, on average 6 years after collection. These are additional examples of efficient rates of new species publications from researchers who undertook both field and herbarium work to describe new tree species.

It is reasonable to interpret that during the 1800s and early 1900s, plants were relatively easier to identify as new species because there were fewer described species to use as reference. After decades of botanical exploration in the Neotropics, a significant amount of plant collections accumulated in the world’s herbaria, meaning that more contemporary botanists likely required more time and effort studying larger collections before determining which plants were actually undescribed species. Furthermore, botanists needed to travel to European and North American herbaria, where the most comprehensive Neotropical plant collections are maintained, to compare and contrast type collections with their own materials and observations before making decisions about potential taxonomic novelties, which may have increased collection to description time lags. In an effort to facilitate access to type specimen images, Macbride visited a number of major European herbaria from 1929 to 1939 and photographed 40,425 types and other historic specimens, representing nearly as many species, chiefly from South America. Macbride’s photographs significantly improved access to type material, and more importantly, recorded images from several type specimens before they were lost during second world war bombing campaigns. These images continue to be a valuable resource available as the Berlin Negatives project at the Field Museum of Natural History (Grimé, 1987). A contemporary effort to enable digital access to type specimens initiated in the early 2000s when the African Plants and Latin American Plants Initiatives were launched to coordinate the digitization of all plant type collections. Those early strategies lead to the Global Plants Initiative, an international partnership of more than 270 herbaria in 70 countries with the goal to digitise, unite, and provide access to type specimens of plants, fungi and algae (Ryan, 2013). To date, over 3 million type specimens are available for viewing, by far the largest online database of biological type specimens. In recent years, major herbaria holding large Neotropical collections including the Netherland’s Naturalis Biodiversity Center, the National Museum of natural History in Paris, the United States National Herbarium, New York Botanical Garden, Royal Botanic Gardens Kew, among others have undertaken the digitization of all of their collections. Furthermore, national herbaria in Latin American countries have initiated massive digitization projects, such as ReFlora virtual herbarium and speciesLink in Brazil (Canhos *et al*., 2022), open data in Mexico (UNAM, 2019) and Bio virtual in Colombia (ICN, 2004). Improved access to digital images from herbarium specimens, coupled with novel analytical approaches based on machine learning and artificial intelligence, have the potential to greatly facilitate taxonomic work (Funk, 2018; Younis *et al*., 2019), leading to more efficient identification and description of new species.

### Publication of new species

With the exception of the monographic work on Guttiferae (Clusiaceae) by Planchon and Triana (1860), the vast majority of new species described during the 1800s until the 1950s originated from field expedition accounts, regional floras, and comprehensive taxonomic treatments that reviewed most of the scientifically known plant species of the time (Figs. 6AG). From the 1960s until the present, most new species of Neotropical trees in our sample were described in taxonomic revisions of single families or genera. Notably, the taxonomic treatment for Chrysobalanaceae (Prance, 1972 b), is the monograph that includes the largest number of newly described species of trees for the region with a total of 90 new taxa. The number of new tree species that have been described in expedition accounts and floristic works vs monographic revisions is comparable, although most modern new species have been described in *ad hoc* publications where a single or few new taxa are described (e. g. Fernandez-Hilario *et al*., 2022; Luján *et al*., 2022; Valdemarin *et al*., 2022). This change in taxonomic publication model may respond to how research productivity is now assessed in most academic institutions. Researchers’ productivity is measured mainly on number of publications and impact factor (e. g. h-index [Hirsch, 2005]) which may be problematic for several reasons (Bi, 2023). Although taxonomic publications rank relatively lower in the impact factor lists, we argue that publishing individual or small numbers of new species should be encouraged and valued, as it is an efficient way to document biodiversity rapidly, which is critical in a world facing large scale biodiversity loss.

### Caveats of this study and final remarks

This study included data from a selection of tree species that are relatively abundant and ecologically important across different Neotropical regions. Time lags between specimen collection and species description may vary for other taxonomic groups or geographical areas. Therefore, our interpretations are limited to particular taxonomic groups and should not be extended further to other components of the Neotropical flora.

We are aware of the challenge to establish a clear causation effect between complex political and social situations of conflict and decline in scientific production. However, we note that largely disruptive events such as war, economic embargo, guerrilla and drug cartelsassociated violence have had far reaching consequences including forced migrations, deep economic crisis, and social unrest in many Latin American countries. Therefore it is only reasonable to assume that the observed slowdown in the pace of new species description can be at least partially influenced by the context of conflict in the region. Governments committed to meet the obligations contracted in international agreements such as the Convention on Biological Diversity (CBD, 1992), including measures towards conservation and sustainable use of biodiversity, need to ensure that researchers can undertake field exploration safely, so species continue to be documented and described at a sensible pace.

Our study shows that the most efficient strategy to describe new species is through collaboration between field botanists collecting in a particular area and researchers with access to natural history collections. Collaborative international research projects should therefore be encouraged and facilitated across scientific and academic institutions so researchers can generate biodiversity data more efficiently. In this context, the UN Biodiversity Conference COP-15 (CBD, 2022) implemented the Kunming-Montreal global biodiversity framework, which includes effective conservation and management of at least 30% of the world’s lands, inland waters, coastal areas and oceans by 2030, with emphasis on areas of particular importance for biodiversity. To this end, one of the four goals adopted by the conference is to establish adequate means of implementation, including financial resources for capacity-building, technical and scientific cooperation, especially in developing countries. Both enhancing international collaboration as well as increasing training opportunities in Latin American countries should be the utmost priorities to accelerate the inventory of the Neotropical biodiversity.

Our results show that one of the most efficient approaches for describing new plant species is by individual researchers undertaking both field collection and species description. Making observations of living plants in the field allows documenting characteristics about the organisms that are inevitably lost during specimen preparation such as colour, scents, 3D shapes, as well as habitat conditions and other sources of valuable biological information. Despite its critical importance, evidence suggests that fieldwork-based investigations have decreased significantly since the 1980s (Rios-Saldaña *et al*., 2014). Here we highlight the importance of continuing and expanding fieldwork and collecting efforts as they provide the fundamental data and materials for biodiversity research and conservation.

We envision the future of describing new species moving towards more integrative approaches where molecular data and digital images will likely play important roles as sources of evidence for species identification. Integration of new species information into larger taxonomic frameworks will largely rely on the willingness of the scientific community to share biodiversity data. Notable advances towards enhancing access to biodiversity data have been accomplished including big data aggregators for species occurrences (e. g. Global Biodiversity Information Facility [GBIF]), repositories for molecular data (e. g. National Center for Biotechnology Information [NCBI]), and historical publications (e. g. Biodiversity Heritage Library [BHL]). Nonetheless, a global open access repository for (new) species descriptions is warranted. Information about taxonomic novelties should be freely accessible to relevant stakeholders including students, conservation practitioners and environmental agencies, particularly from low-income tropical countries, where this information has the greatest potential to inform and influence urgently needed conservation efforts.

## Acknowledgements

ML is thankful to Dr. Eimear Nic Lughadha for encouraging the preparation of this manuscript and providing valuable advice on the original idea. Three anonymous reviewers provided helpful comments that significantly improved early versions of the manuscript. The authors humbly dedicate this collective effort to honour the memory Dr. Jerzy Rzedowski, a heartfelt tribute to our esteemed co-author whose contributions to the Neotropical botany will be everlasting.

## References

Antonelli, A. & Sanmartín, I. (2011). Why are there so many plant species in the Neotropics?. Taxon, 60 (2), 403–414. DOI: 10.1002/tax.602010

Archer, W.A. (1962). Adolpho Ducke, botanist of the Brazilian Amazon (1876-1959). Taxon, 11(8), 233–242. DOI: 10.2307/1217031.

Baber, Z. (2016). The Plants of Empire: Botanic Gardens, Colonial Power and Botanical Knowledge. Journal of Contemporary Asia, 46 (4), 659–679. DOI: 10.1080/00472336.2016.1185796.

Bebber, D.P., Wood, J.R., Barker, C. & Scotland, R.W. (2014). Author inflation masks global capacity for species discovery in flowering plants. New Phytologist, 201(2), 700–706. DOI: 10.1111/nph.12522.

Bedoya, A.M. & Olmstead R.G. (2022). Strange but common in isolated environments: new records of Marathrum (Podostemaceae) in rivers of Colombia. Aquatic Botany, 177: 103483. DOI: 10.1016/j.aquabot.2021.103483.

Beentje, H. (2020). The Kew Plant Glossary an illustrated dictionary of plant terms (2nd ed.). Royal Botanical Gardens, Kew.

Bi, H.H. (2023). Four problems of the h-index for assessing the research productivity and impact of individual authors. Scientometrics, 128, 2677–2691. 10.1007/s11192-022-04323-8.

Botanic Gardens Conservation International. (2021). State of The World’s Trees Report. Global Tree Portal. https://www.bgci.org/resources/bgci-databases/globaltree-portal/ (Accessed on 8/11/2022). DOI: 10.1016/j.aquabot.2021.103483.

Boyd, I.L., Freer-Smith, P.H., Gilligan, C.A. and Godfray, H.C.J. (2013). The consequence of tree pests and diseases for ecosystem services. Science, 342 (6160), 1235773. DOI: 10.1126/science.1235773.

Canhos, D.A.L., Almeida, E.A., Assad, A.L., Cunha Bustamante, M.M.D., Canhos, V.P., Chapman, A.D., Giovanni, R.D., Imperatriz-Fonseca, V.L., Lohmann, L.G., Maia, L.C. and Miller, J.T. (2022). speciesLink: rich data and novel tools for digital assessments of biodiversity. Biota Neotropica, 22(spe), e20221394. DOI: 10.1590/1676-0611BN-2022-1394.

Convention on Biological Diversity (CBD). (1992). United Nations Environmental Programme. Secretariat of the Convention on Biological Diversity. https://www.cbd.int/doc/legal/cbd-en.pdf. Accessed 16 February 2023.

Convention on Biological Diversity (CBD). (2022). Conference of the Parties to the Convention on Biological Diversity. Capacity-building and development and technical and scientific cooperation. https://www.cbd.int/doc/c/f071/ba75/4aeaaa842acdaf622d1b6a18/cop15-l-28-en.pdf. Accessed 16 February 2023.

Cowan, R.S. (1967). *Swartzia* (Leguminosae, Caesalpinioideae Swartzieae). Flora Neotropica, 1, 1–228. DOI: https://www.jstor.org/stable/4393651.

Cuatrecasas, J. (1950). Notas a la Flora de Colombia, X, Guttiferae. Revista de la Academia Colombiana Ciencias Exactas 8, 33–64.

De Queiroz K. (2007). Species concepts and species delimitation. Systematic Biology. 56 (6), 879–886.

Engler, A. (1888). Guttiferae and Quiinaceae. Pp. 382–486 in: Urban, I. (ed.) Flora Brasiliensis, vol. 12,1. Leipzig: Fleischer.

Fernandez-Hilario, R., Gonzáles, R.D.P.R., Villanueva-Espinoza, R., Lajo, L., Sato, A.A.W., Fontaine, B., Perrard, A. & Bouchet, P. (2012). 21 years of shelf life between discovery and description of new species. Current Biology, 22 (22), R943–R944. DOI: 10.1016/j.cub.2012.10.029.

Freitas, C.G., Bacon, C.D., Souza-Neto, A.C. & Collevatti, R.G. (2019). Adjacency and area explain species bioregional shifts in neotropical palms. Frontiers in Plant Science, 10, 55. DOI: 10.3389/fpls.2019.00055.

Fritsch, C. (1889). Beiträge zur Kenntniss der Chrysobalanaceen. I: Conspectus generis Licaniae. Annalen des Kaiserlich-Königlichen Naturhistorischen Hofmuseums 4, 33–60.

Funk, V.A. (2018). Collections based science in the 21st century. Journal of Systematics and Evolution, 56 (3), 175–193. DOI: 10.1111/jse.12315

Gentry, A.H. (1973). Bignoniaceae in Flora of Panama. Annals Missouri Botanical Garden 60(3), 781–977. DOI: 10.2307/2395140

Govaerts, R. (2001). How many species of seed plants are there? Taxon 50: 1085–1090. DOI: 10.2307/1224723.

Govaerts, R., Nic Lughadha, E., Black, N., Turner R. & Paton A. (2021). The World Checklist of Vascular Plants, a continuously updated resource for exploring global plant diversity. Scientific Data 8 (1), 215. DOI: 10.1038/s41597-021-00997-6.

Goodwin, Z.A., Muñoz-Rodríguez, P., Harris, D.J., Wells, T., Wood, J.R., Filer, D. & Scotland, R.W. (2020) How long does it take to discover a species?. Systematics and Biodiversity, 18 (8), 784–793. DOI: 10.1080/14772000.2020.1751339.

Grillo, I. (2013). Mexican Cartels: A Century of Defying U.S. Drug Policy. The Brown Journal of World Affairs, 20 (1), 253–265.

Grimé, W.E. (1987). Type photographs at the Field Museum of Natural History. Taxon, 36(2), 425–426. DOI: 10.2307/1221436.

Grisebach, A.H.R. (1866). Catalogus plantarum Cubensium exhibens collectionem Wrightianam aliasque minores ex insula Cuba missas. Lipsiae apud Guilielmum Engelmann.

Hiepko, P. (1987). The collections of the Botanical Museum Berlin-Dahlem (B) and their history. Englera, 7, 219–252.

Hirsch, J. E. (2005). An index to quantify an individual’s scientific research output. Proceedings of the National Academy of Sciences of the United States of America, 102 (46), 16569–16572. DOI: 10.1073/pnas.0507655102.

Howard, R.A. (1983). The plates of Aublet’s Histoire des Plantes de la Guiane Francoise. Journal of the Arnold Arboretum, 64 (2), 255–292. DOI: https://www.jstor.org/stable/43782107

Huber, O. (1996). José Cuatrecasas. Acta Botánica Venezuélica, 19 (1), 76–81. DOI: https://www.jstor.org/stable/41740533

Instituto de Ciencias Naturales (ICN), Facultad de Ciencias, Universidad Nacional de Colombia. (2004) onwards. Colecciones en Línea. Available: http://www.biovirtual.unal.edu.co. Accessed on 06 Feb 2023.

Irwin, A. (2023). The race to understand Colombia’s exceptional biodiversity. Nature, 619, 450–453. https://www.nature.com/articles/d41586-023-02300-6.pdf.

Joppa, L.N., Roberts, D.L., Myers, N. & Pimm, S.L. (2011). Biodiversity hotspots house most undiscovered plant species. Proceedings of the National Academy of Sciences, 108 (32), 13171–13176. DOI: 10.1073/pnas.1109389108

LeGrand, C.C. (2003). The Colombian crisis in historical perspective. Canadian journal of Latin American and Caribbean studies, 28 (55/56), 165–209. DOI: 10.1080/08263663.2003.10816840.

Li, F. W. (2021). Decolonizing botanical genomics. Nature Plants, 7 (12), 1542–1543. 10.1038/s41477-021-01041-6.

Luján, M., Cacho, N.I., Pérez-Farrera, M.Á. & Hammel, B. (2021). *Clusia falcata* (Clusiaceae), an endangered species with exceptionally narrow leaves endemic to Chiapas, Mexico. Kew Bulletin 76, 645–650. DOI: 10.1007/S12225-021-09988-7.

Mishler B. D., & Brandon R. N. (1987). Individuality, pluralism, and the phylogenetic species concept. Biology and Philosophy, 2, 397–414. DOI: 10.1007/BF00127698.

Mishler, B. D. & Donoghue M. J. (1982). Species concepts: a case for pluralism. Systematic Zoology, 31 (4), 491–503. DOI: 10.2307/2413371.

Mishler, B. D. (2021). What, if anything, are species? CRC Press, Boca Raton.

Nagoya Protocol on Access to Genetic Resources and the Fair and Equitable Sharing of Benefits Arising from their Utilisation to the Convention on Biological Diversity: text and annex (2011). Secretariat of the Convention on Biological Diversity. United Nations Environmental Programme. Montreal. 15 pp. https://www.cbd.int/abs/doc/protocol/nagoyaprotocol-en.pdf. Accessed 16 February 2023.

Lucas, E.J., Haigh, A.L., Castellanos, C., Aguilar-Cano, J., Biggs, N., Castellanos, C.C., Fabriani, F., Frisby, S., García, L., Klitgård, B.B. & Morales-Puentes, M.E. (2023). An updated checklist of Araceae, Leguminosae and Myrtaceae of the department of Boyacá, Colombia, including keys to genera and new occurrence records. Phytotaxa, 589 (2), 137–178. DOI: 10.11646/phytotaxa.589.2.4.

Lughadha, E.N., Govaerts, R., Belyaeva, I., Black, N., Lindon, H., Allkin, R., Magill, R.E. & Nicolson, N. (2016). Counting counts: revised estimates of numbers of accepted species of flowering plants, seed plants, vascular plants and land plants with a review of other recent estimates. Phytotaxa, 272 (1), 82–88. DOI: 10.11646/phytotaxa.272.1.5

Olson, D.M., Dinerstein, E., Wikramanayake, E.D., Burgess, N.D., Powell, G.V., Underwood, E.C., D’amico, J.A., Itoua, I., Strand, H.E., Morrison, J.C. and Loucks, C.J., (2001). Terrestrial Ecoregions of the World: A New Map of Life on Earth: A new global map of terrestrial ecoregions provides an innovative tool for conserving biodiversity. BioScience, 51 (11), 933–938. DOI: 10.1641/00063568(2001)051[0933:TEOTWA]2.0.CO;2.

Planchon, J.E. & J. Triana. (1860). Mémoire sur la famille de Guttifères. Annales des Sciences Naturelles. Botanique Serie IV. 14: 226–367.

Prance G. T. (1972 a). An index of plant collectors in Brazilian Amazonia. Acta Amazonica 1 (1), 25–65, DOI: 10.1590/1809-43921971011025.

Prance G. T. (1972 b). Monograph of Chrysobalanaceae. Flora Neotropica Monographs 9: 1–406. DOI: http://www.jstor.org/stable/4393681.

Raven, P.H., Gereau, R.E., Phillipson, P.B., Chatelain, C., Jenkins, C.N. and Ulloa Ulloa, C., (2020). The distribution of biodiversity richness in the tropics. Science Advances, 6 (37), p.eabc6228. DOI: 10.1126/sciadv.abc6228.

Ríos-Saldaña, C. A., Delibes-Mateos, M., & Ferreira, C. C. (2018). Are fieldwork studies being relegated to second place in conservation science?. Global ecology and conservation, 14, e00389.

Ronda-Pupo G.A. (2021). Cuba-U.S. scientific collaboration: Beyond the embargo. PLoS One 16 (7): e0255106. DOI: 10.1371/journal.pone.0255106.

Ryan, D. (2013). The Global Plants Initiative celebrates its achievements and plans for the future. Taxon, 62 (2), 417–418. DOI: https://www.jstor.org/stable/taxon.62.2.417

Rzedowski, J. (1967). Faustino Miranda, 1905–1964. Brittonia, 19 (1), 95–98.

Rzedowski, J. (1981). A century of botany in Mexico. Botanical Sciences, 40, 1–14. DOI: 10.17129/botsci.1183.

Schomburgk, O.A. (1931). Robert Hermann Schomburgk’s Travels in Guiana and on the Orinoco During the Years 1835–1839. The Argosy Company Ltd. Georgetown.

Seaward, M.R. (2010). Richard E. Schultes and the botanist-explorer Richard Spruce (1817–1893). Harvard Papers in Botany 15 (2), 447–454. DOI: https://www.jstor.org/stable/41761709.

Sokal, R.R. & Crovello T. (1970). The biological species concept: A critical evaluation. American Naturalist 104 (936), 127–153. DOI: https://www.jstor.org/stable/2459191

Standley, P.C. (1931). Erik L. Ekman. Science, 73, 255–256. DOI: 10.1126/science.73.1888.255.

Stearn, W.T. (1965). Grisebach’s flora of the British West Indian Islands: A biographical and bibliographical introduction. Journal of the Arnold Arboretum, 46 (3), 243–285.

Taylor R.W. (1983). Descriptive taxonomy: past, present, and future. In Australian Systematic entomology: a bicentenary perspective, Highley E, Taylor RW (eds). CSIRO: Melbourne.

Universidad Autonoma de Mexico (UNAM). (2019) onwards. Portal de datos abiertos. Colecciones Universitarias. Available: https://datosabiertos.unam.mx/. Accessed on 06 Feb 2023.

Valdemarin, K.S., Camargo, P.H., Moreno, D.J., Souza, V.C., Lucas, E. and Mazine, F.F., (2022). *Eugenia paranapanemensis* (Myrtaceae), the Pitanga-amarela, and a Key to *Eugenia* sect. *Eugenia* Species from São Paulo State, Brazil. Systematic Botany, 47 (2), 498–505. DOI: 10.1600/036364422X16512564801669.

Wade, L. (2018). Colombian scientists race to study once-forbidden territory before it is lost to development or new conflict. Science. DOI:10.1126/science.aat9905.

Wells, T., Carruthers, T., Muñoz-Rodríguez, P., Sumadijaya, A., Wood, J.R. and Scotland, R.W. (2022). Species as a heuristic: reconciling theory and practice. Systematic Biology, 71(5), 1233–1243. DOI: 10.1093/sysbio/syab087.

Younis S., Weiland C., Hoehndorf R., Dressler S., Hickler T., Seeger B., Schmidt M. (2018). Taxon and trait recognition from digitized herbarium specimens using deep convolutional neural networks. Botany Letters, 165 (3-4), 377–383. DOI: 10.1080/23818107.2018.1446357.

